# Incomplete inhibition of HIV infection results in more HIV infected lymph node cells by reducing cell death

**DOI:** 10.1101/163352

**Authors:** Laurelle Jackson, Jessica Hunter, Sandile Cele, Isabella Markham Ferreira, Andrew Young, Farina Karim, Rajhmun Madansein, Kaylesh J. Dullabh, Chih-Yuan Chen, Noel J. Buckels, Yashica Ganga, Khadija Khan, Mikaël Boullé, Gila Lustig, Richard A. Neher, Alex Sigal

## Abstract

HIV has been reported to be cytotoxic *in vitro* and in lymph node infection models. Using a computational approach, we found that partial inhibition of transmission which involves multiple virions per cell could lead to increased numbers of live infected cells if the number of viral DNA copies remains above one after inhibition, as eliminating the surplus viral copies reduces cell death. Using a cell line, we observed increased numbers of live infected cells when infection was partially inhibited with the antiretroviral efavirenz or neutralizing antibody. We then used efavirenz at concentrations reported in lymph nodes to inhibit lymph node infection by partially resistant HIV mutants. We observed more live infected lymph node cells, but with fewer HIV DNA copies per cell, relative to no drug. Hence, counterintuitively, limited attenuation of HIV transmission per cell may increase live infected cell numbers in environments where the force of infection is high.

## Introduction

HIV infection is known to result in extensive T cell depletion in lymph node environments (Sanchez et al, 2015), where infection is most robust (Brenchley et al, 2004; Doitsh et al, 2010; Doitsh et al, 2014; Finkel et al, 1995; Galloway et al, 2015; Mattapallil et al, 2005). Depletion of HIV infectable target cells, in addition to onset of immune control, is thought to account for the decreased replication ratio of HIV from an initial peak in early infection (Bonhoeffer et al, 1997; Nowak & May, 2000; Perelson, 2002; Phillips, 1996; Quinones-Mateu & Arts, 2006; Ribeiro et al, 2010; Wodarz & Levy, 2007). This is consistent with observations that individuals are most infectious in the initial, acute stage of infection, where the target cell population is relatively intact and produces high viral loads (Hollingsworth et al, 2008; Wawer et al, 2005).

T cell death occurs by several mechanisms, which are either directly or indirectly mediated by HIV infection. Accumulation of incompletely reverse transcribed HIV transcripts is sensed by interferon-γ –inducible protein 16 (Monroe et al, 2014) and leads to pyroptotic death of incompletely infected cells by initiating a cellular defence program involving the activation of caspase 1 (Doitsh et al, 2010; Doitsh et al, 2014; Galloway et al, 2015). HIV proteins Tat and Env have also been shown to lead to cell death of infected cells through CD95 mediated apoptosis following T cell activation (Banda et al, 1992; Westendorp et al, 1995a; Westendorp et al, 1995b). Using SIV infection, it has been shown that damage to lymph nodes due to chronic immune activation leads an environment less conducive to T cell survival (Zeng et al, 2012). Finally, double strand breaks in the host DNA caused by integration of the reverse transcribed virus results in cell death by the DNA-PK mediated activation of the p53 response (Cooper et al, 2013).

The lymph node environment is conducive to HIV infection due to the presence of infectable cells (Deleage et al, 2016; Embretson et al, 1993; Tenner-Racz et al, 1998), the proximity of cells to each other and lack of flow which should enable cell-to-cell HIV spread (Baxter et al, 2014; Dale et al, 2011; Groot et al, 2008; Groppelli et al, 2015; Gummuluru et al, 2002; Hubner et al, 2009; Jolly et al, 2004; Jolly et al, 2011; Munch et al, 2007; Sherer et al, 2007; Sourisseau et al, 2007; Sowinski et al, 2008), and decreased penetration of antiretroviral therapy (ART) (Fletcher et al, 2014a). Multiple infections per cell have been reported in cell-to-cell spread of HIV (Baxter et al, 2014; Boullé et al, 2016; Dang et al, 2004; Del Portillo et al, 2011; Dixit & Perelson, 2004; Duncan et al, 2013; Law et al, 2016; Reh et al, 2015; Russell et al, 2013; Sigal et al, 2011; Zhong et al, 2013), where an interaction between the infected donor cell and the uninfected target results in directed transmission of large numbers of virions (Baxter et al, 2014; Groppelli et al, 2015; Hubner et al, 2009; Sowinski et al, 2008). This is in contrast to cell-free infection, where free-floating virus finds target cells through diffusion. Both modes occur simultaneously when infected donor cells are cocultured with targets, though the cell-to-cell route is the main cause of multiple infections per cell (Hubner et al, 2009). In the lymph nodes, several studies showed multiple infections (Gratton et al, 2000; Jung et al, 2002; Law et al, 2016) while another study did not (Josefsson et al, 2013). One explanation for the divergent results is that different cell subsets are infected to different degrees. For example, T cells were shown not to be multiply infected in the peripheral blood compartment (Josefsson et al, 2011). However, more recent work investigating markers associated with HIV latency in the face of ART found that the average number of HIV DNA copies per cell is greater than one in 3 out of 12 individuals in the face of ART in the CD3 positive, CD32a high CD4 T cell subset (Descours et al, 2017). In the absence of suppressive ART, it would be expected that the number of HIV DNA copies per cell would be higher.

Multiple viral integration attempts per cell may increase the probability of death. One consequence of HIV mediated death may be that attenuation of infection may increase viral replication by increasing the number of live targets. Indeed, it has been suggested that more attenuated HIV strains result in more successful infections in terms of the ability of the virus to replicate in the infected individual (Arien et al, 2005; Nowak & May, 2000; Payne et al, 2014; Quinones-Mateu & Arts, 2006; Wodarz & Levy, 2007).

Here we experimentally examined the effect of attenuating cell-to-cell spread by using HIV inhibitors. We observed that partially inhibiting infection with drug or antibody resulted in an increase in the number of live infected lymph node cells, experimentally demonstrating for the first time at the cellular level that attenuation of HIV infection can result in an increase in live infected cells under specific infection conditions.

## Results

We introduce a model of infection where each donor to target transmission leads to an infection probability *r* and death probability *q*. The probability of infection of a target given *λ* infecting virions is *1-(1-r)*^*λ*^ (Sigal et al, 2011). The probability of a cell to be infected and not die after it has been exposed to *λ* transmitted virions is therefore:

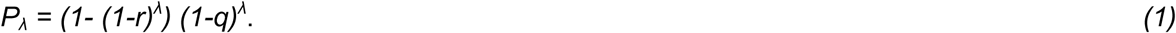

This model makes several simplifying assumptions: 1) all transmissions have equal probabilities to infect targets. 2) The probability for a cell to die from each transmission is equal between transmissions. 3) Productive infection and death are independent events.

Since antiretroviral drugs lead to a reduction in the number of infecting virions by, for example, decreasing the probability of reverse transcription in the case of reverse transcriptase inhibitors, we introduced a drug strength value *d*, where *d=1* in the absence of drug and *d>1* in the presence of drug. In the presence of drug, *λ* is decreased to **λ**/*d*. The drug therefore tunes *λ*, and if the antiretroviral regimen is fully suppressive, *λ/d* is expected to be below what is required for ongoing replication. The probability of a cell to be infected and live given drug strength *d* is therefore:

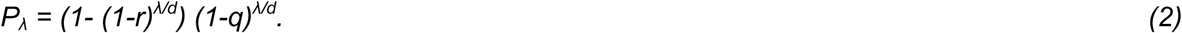

By this model, the probability to be infected and not die *P*_*λ*_ depends on the number of virions and has two parameters, *r* and *q*, as well as two measurable inputs, *λ* and *d*, where *d/λ* measures the effect of drug relative to infection strength per cell. Analysis of the probability of a cell to survive and be infected as a function of *r* and *q* shows that at each drug strength *d/*λ, *P*_*λ*_ increases as the probability of infection *r* increases, regardless of the probability of death per infection *q* (Figure 1). Whether and how *P*_*λ*_ behaves when drug strength *d/λ* increases depends on parameter values of *r* and *q*. A subset of parameter values results in a peak in the number of infected cells at intermediate *d/λ*, decreasing as drug strength increases further (Figure 1). We refer to such a peak in infected numbers as an infection optimum. As *q* increases, the cost of multiple infections per cell increases, and the infection optimum shifts to higher *d/λ* values. A fall from the infection optimum at decreasing *d/λ* is driven by increasing cell death as a result of increasing infection attempts per cell. This slope is therefore shallower, and peaks broader, at low *q* values (Figure 1).

**Figure 1:**
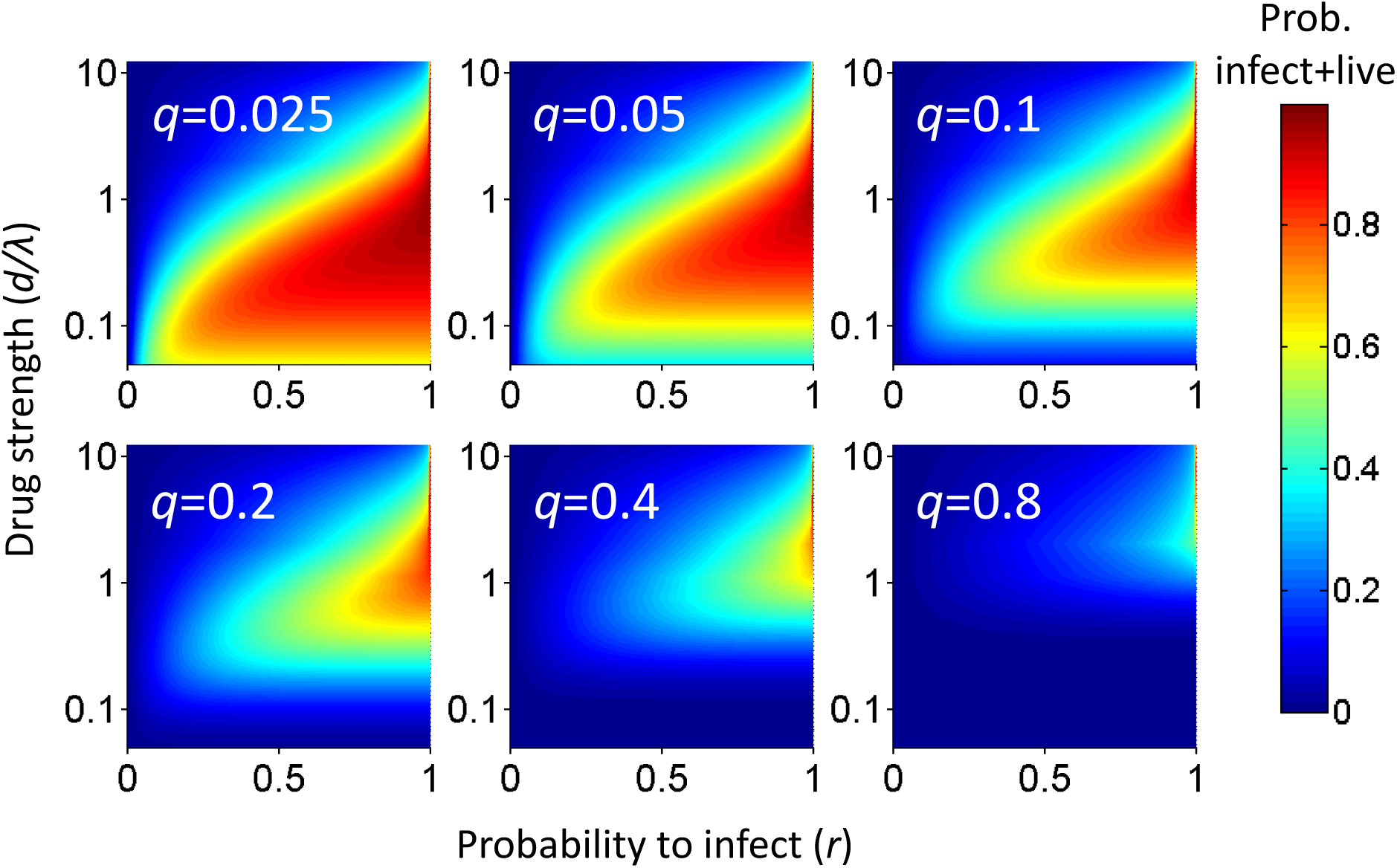
Probability for a cell to be infected and live as a function of inhibitor. Probability for a cell to be infected and live was calculated for 20 infection attempts (*λ*) and represented as a heat map. Drug strength as *d/λ* is on the y-axis, and the probability per virion to infect (*r*) is on the x-axis. Each plot is the calculation for one value of the probability per virus to die (*q)* denoted in white in the top left corner.

Given that an infection optimum is dependent on parameter values, we next tested whether these parameter values occur experimentally in HIV infection. We therefore first tested for an infection optimum in the RevCEM cell line engineered to express GFP upon HIV Rev protein expression (Wu et al, 2007). We subcloned the cell line to obtain clone E7 which maximized the frequency of GFP positive cells upon infection (Boullé et al, 2016).

We needed to detect the number of integration attempts per cell *λ*. To estimate this, we used PCR to detect the number of reverse transcribed copies of viral DNA in the cell by splitting each individual infected cell over multiple wells and detecting the number of wells with the expected PCR product. Hence, the number of positive wells indicates the minimum number of viral DNA copies per cell, since more than one copy can be contained within the same well (Josefsson et al, 2011; Josefsson et al, 2013). We first examined the number of viral DNA copies in ACH-2 cells, reported to contain single inactive HIV integration per genome (Chun et al, 1997; O’Doherty et al, 2002). We sorted individual ACH-2 cells and split the single cell lysates into multiple wells (Figure 2 – figure supplement 1A). About one quarter of cells showed a PCR product of the expected size (Figure 2 – figure supplement 1B). Similar results were obtained with cell-free infection of E7 RevCEM cells (Figure 2 – figure supplement 1C).

**Figure 2:**
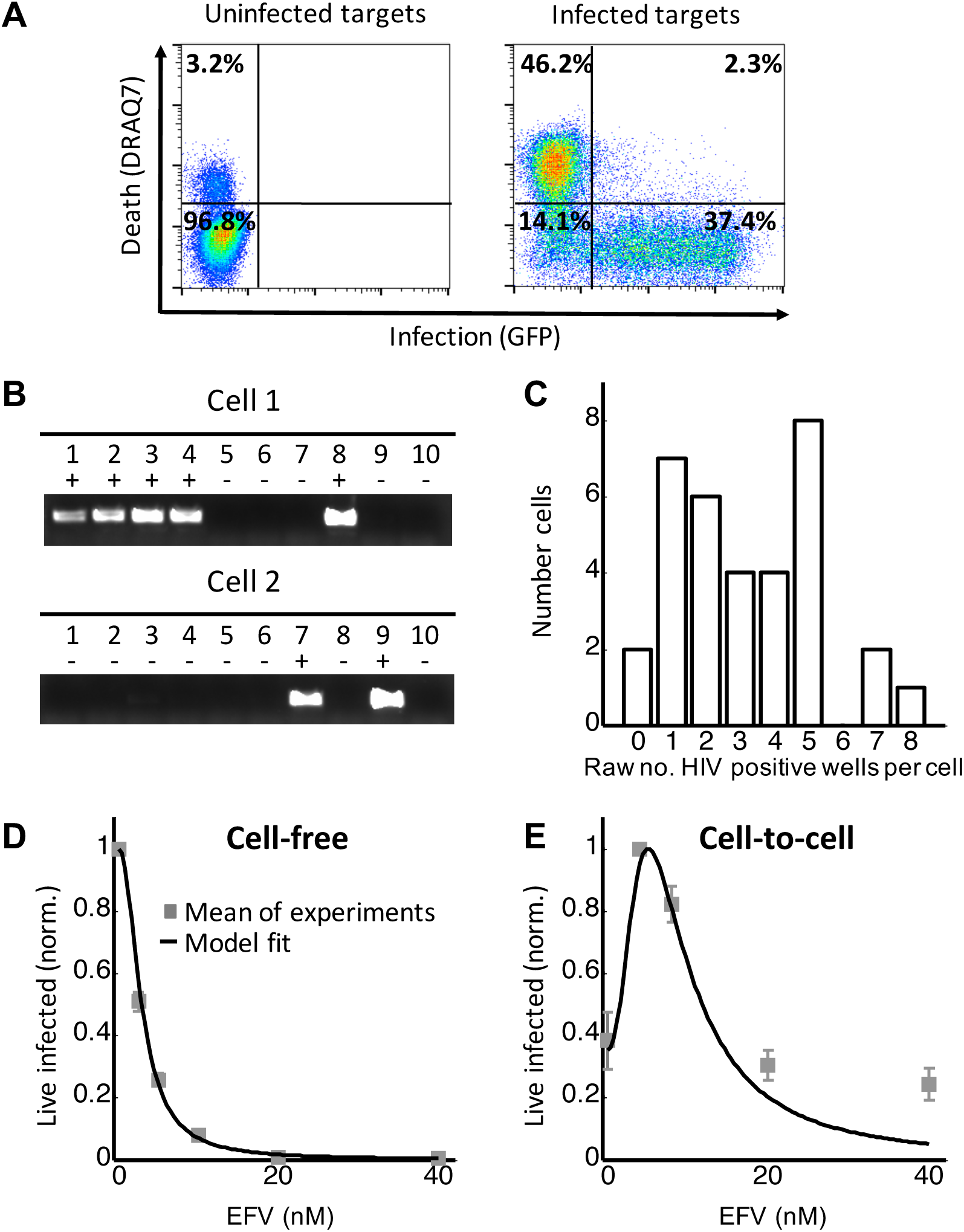
Partial inhibition of HIV in coculture infection increases the number of live infected cells. (A) Experimental system to detect live infected cells by flow cytometry. Left plot shows uninfected E7 reporter cells, and right plot shows E7 cells after 2 days of infection. Infected cells are detected by HIV dependent GFP reporter expression (x-axis), while dead cells as detected by the intracellular presence of the death indicator dye DRAQ7 (y-axis). Live infected cells are in the bottom right quadrant. (B)To quantify HIV DNA copy number per cell, GFP positive cells were sorted into individual wells of a multi-well plate at 1 cell per well and lysed. Each lysed cell was further subdivided into 10 wells and nested PCR performed to detect whether the well contained a copy of HIV DNA. The sum of PCR positive wells for a given cell was used as the raw HIV DNA copy number for that cell. (C) Histogram of the raw number of detected HIV DNA copies per cell. Results are of 34 cells from 3 independent experiments (n=4 for experiment 1, n=20 for experiment 2, n=10 for experiment 3). (D) The number of live infected cells normalized by the maximum number of live infected cells in cell-free infection as a function of EFV. Black line represents best-fit model for EFV suppression of single virion infection, with IC_50_=2.9nM and Hill coefficient of 2.1. Shown are means and standard errors for 3 independent experiments, with a minimum of 50,000 cells collected per data point per experiment by flow cytometry. (E) The number of live infected cells normalized by the maximum number of live infected cells in coculture infection as a function of EFV. Black line represents best-fit model for the effect of EFV on infection with multiple virions according to Equation (2), with drug strength (*d*) values calculated based on the cell-free infection data. The fits recapitulated the experimental results when *r=*0.4 and *q=*0.11. Shown are means and standard errors for 3 independent experiments, with a minimum of 50,000 cells collected per data point per experiment by flow cytometry.

To investigate the effect of multiple infection attempts per cell, we used coculture infection, where infected (donor) cells are co-incubated with uninfected (target) cells and lead to cell-to-cell spread. Live infected target cells were assayed after 2 days in coculture with infected donors. Live infected cells were identified based on the absence of cell death indicator dye DRAQ7 fluorescence, and presence of GFP (Figure 2A). We used approximately 2% infected donor cells as our input, and detected the number of HIV DNA copies per cell by sorting individual cell and splitting the individual cell lysates over 10 wells of an agarose gel (Figure 2B). We obtained a wide distribution of viral DNA copies per cell, ranging from 0 to 8 (Figure 2C). We computationally corrected the detected number of DNA copies for the sensitivity of our PCR reaction as determined by the ACH-2 results (Materials and Methods). On average we obtained 14 copies per cell after correction.

To tune *λ*, we added the HIV reverse transcriptase inhibitor efavirenz (EFV) to infections. To calculate *d*, we used cell-free infection, which we verified results in single integrations per cell (Figure 2D, see Figure 2–figure supplement 2 for logarithmic y-axis plot). For cell-free infection, we approximate *d = 1/Tx,* where *Tx* is defined as the number of infected cells with drug divided by the number of infected cells without drug (see Materials and Methods and (Sigal et al, 2011)), equivalent to the fraction of unaffected cells in a previously described approach modelling the infection response to inhibitor with two parameters: the 50% inhibitory drug concentration (IC_50_) and the Hill coefficient for drug action (Shen et al, 2008). We fitted the observed response of infection to EFV to using this approach to estimate *d* across a range of EFV concentrations. The best fit of the model to the cell-free data using EFV sensitive HIV showed a monotonic decrease with IC_50_=2.9nM and Hill coefficient of 2.1 (Figure 2D, black line).

We next dialled in EFV to tune *λ/d* in coculture infection. We observed a peak in the number of live infected target cells at 4nM EFV (Figure 2E). Equation (2) best fit the behaviour of infection when *r =*0.4 *and q =*0.11 (Figure 2E, black line). Hence an infection optimum is present in the cell line infection system.

We repeated the experiment with EFV resistant HIV. To derive the resistant mutant, we cultured EFV sensitive HIV in our reporter cell line in the presence of EFV. We obtained the L100I partially resistant mutant. We then replaced the reverse transcriptase of the wild type molecular clone with the mutant reverse transcriptase gene (Materials and Methods). We derived *d*_*mut*_ for the L100I mutant using cell-free mutant infection (Figure 3A, see Figure 3–figure supplement 1 for logarithmic y-axis plot). L100I mutant was found to have an IC_50_=29nM EFV and Hill coefficient of 2.0 (Figure 3A, black line).

**Figure 3:**
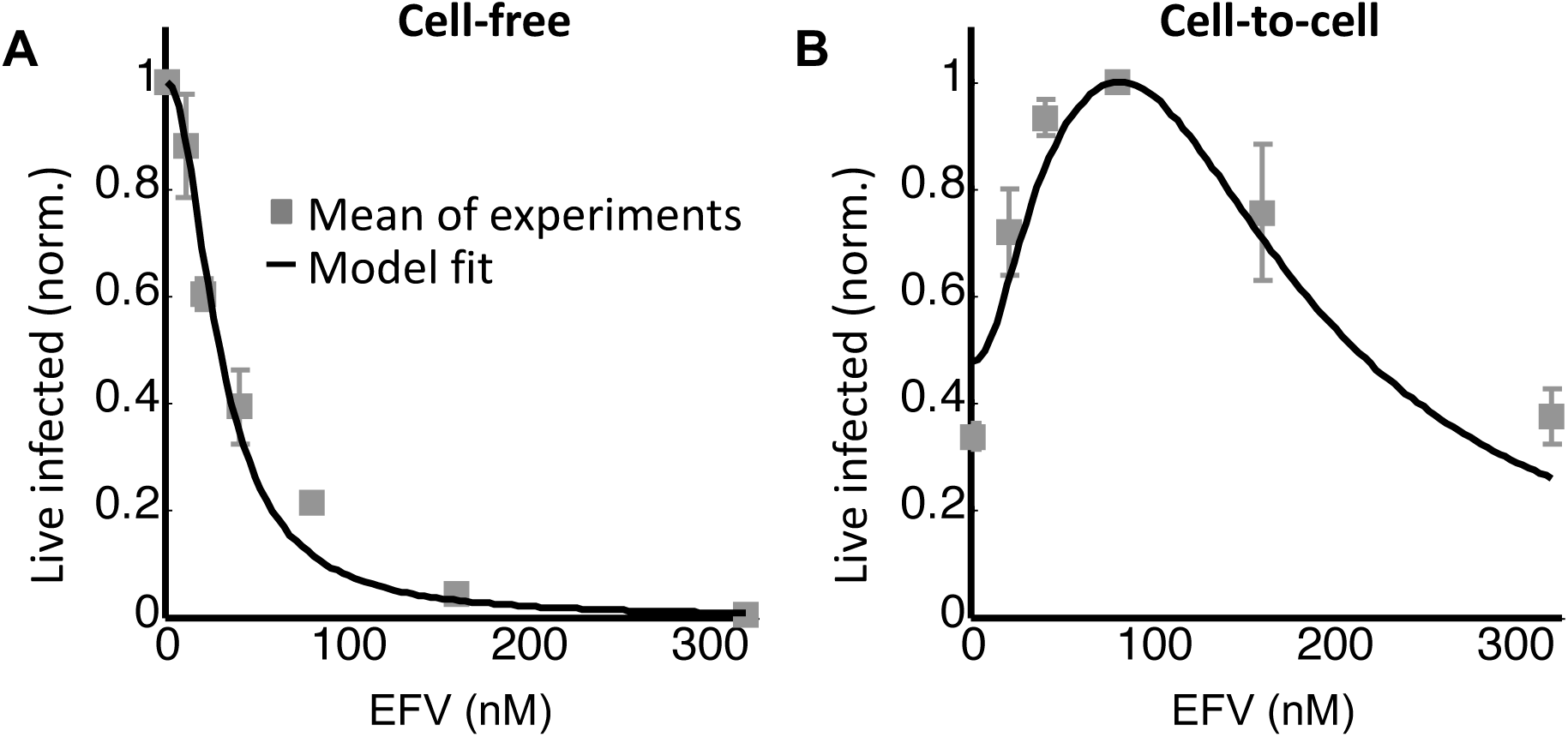
Partial inhibition of the EFV resistant L100I mutant shifts the peak of live infected cells to higher EFV concentrations. (A) The number of live infected cells normalized by the maximum number of live infected cells in cell-free infection as a function of EFV for the L100I mutant. Black line represents best-fit model for EFV suppression of single virion infection, with IC_50_=29nM and Hill coefficient of 2.0. Shown are means and standard errors for 3 independent experiments, with a minimum of 50,000 cells collected per data point per experiment by flow cytometry. (B) The number of live infected cells normalized by the maximum number of live infected cells in coculture infection by the L100I mutant as a function of EFV. Black line represents best-fit model for the effect of EFV on infection with multiple virions according to Equation (2), with *d* values calculated based on the cell-free infection data for the L100I mutant. The fits recapitulated the experimental results when *r*=0.9 and *q*=0.06. Shown are means and standard errors for 3 independent experiments, with a minimum of 50,000 cells collected per data point per experiment by flow cytometry.

We next performed coculture infection. Like with wild type HIV coculture infection, there was a peak in the number of live infected target cells for the L100I mutant infection, but it was shifted to 80nM EFV (Figure 3B), an EFV concentration which a decreased cell-free mutant infection approximately 9-fold and *λ/d*_*mut*_ *∼* 2. Fits were obtained to Equation (2) using *d*_*mut*_ values and *λ* measured for wild type infection. The fits recapitulated the experimental results when *r =*0.9 and *q =*0.06 (Figure 3B, black line).

In order to examine whether a peak in live infected targets can be obtained with an unrelated inhibitor, we used the HIV neutralizing antibody b12. This antibody is effective against cell-to-cell spread of HIV (Baxter et al, 2014; Reh et al, 2015). We obtained a peak with 5ug/ml b12 (Figure 4). The b12 concentration that resulted in a peak number of live infected cells was the same for wild type virus and the L100I mutant, showing L100I mutant fitness gain was EFV specific.

**Figure 4:**
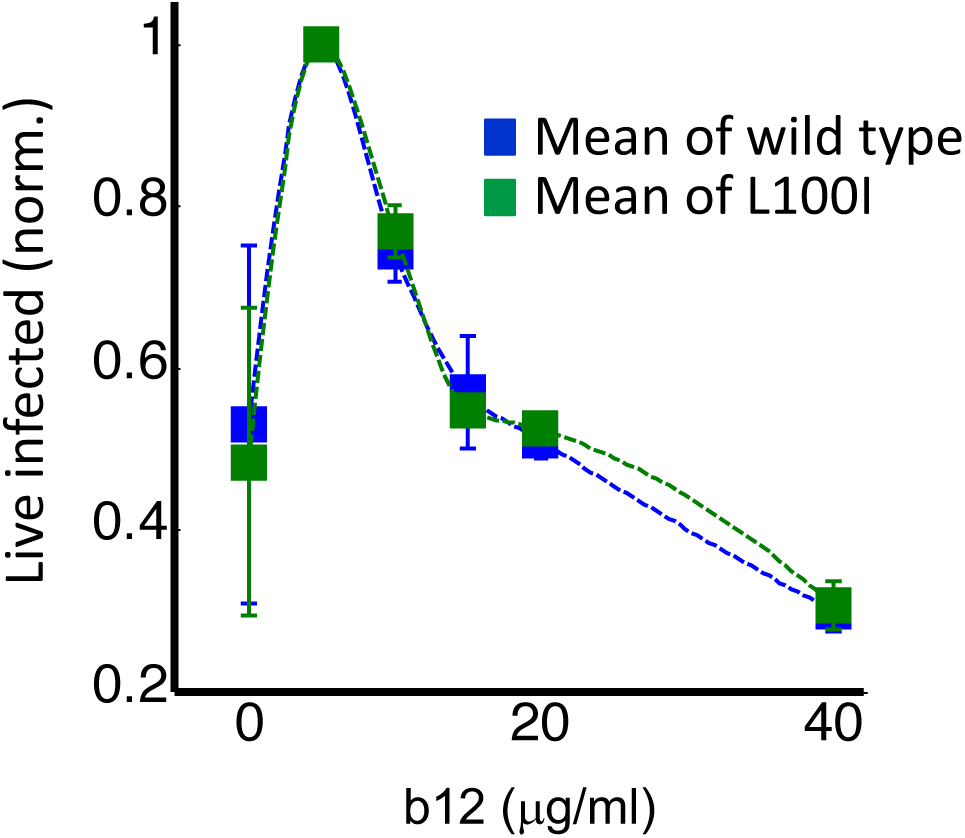
Partial inhibition of coculture infection with neutralizing antibody results in higher numbers of live infected cells. Shown are the numbers of live infected cells normalized by the maximum number of live infected cells in coculture infection as a function of b12 antibody concentration. Infection was by either EFV sensitive HIV (blue) or the L100I EFV resistant mutant (green). Dotted lines are a guide to the eye. Shown are means and standard errors for 3 independent experiments, with a minimum of 50,000 cells collected per data point per experiment by flow cytometry.

While the RevCEM cell line is a useful tool to illustrate the principles governing the formation of an infection optimum, the sensitivity of such an optimum to parameter values would make its presence in primary HIV target cells speculative. We therefore investigated whether a fitness optimum occurs in primary human lymph node cells, the anatomical site of which would be most likely associated with a high force of infection. We derived human lymph nodes from HIV negative individuals from indicated lung resections (S Table 1), cellularized the lymph node tissue, and infected the resulting lymph node cells with HIV by coculture infection. A fraction of the cells was infected by cell-free and used as infected donor cells. We added these to uninfected target cells from the same lymph node, and detected the number of live infected cells after 4 days of infection with the L100I EFV resistant mutant in the face of EFV. We detected the number of live infected cells by the exclusion of dead cells with the death detection dye eFluor660 followed by single cell staining for HIV Gag using anti-p24 antibody (Figure 5A).

**Figure 5:**
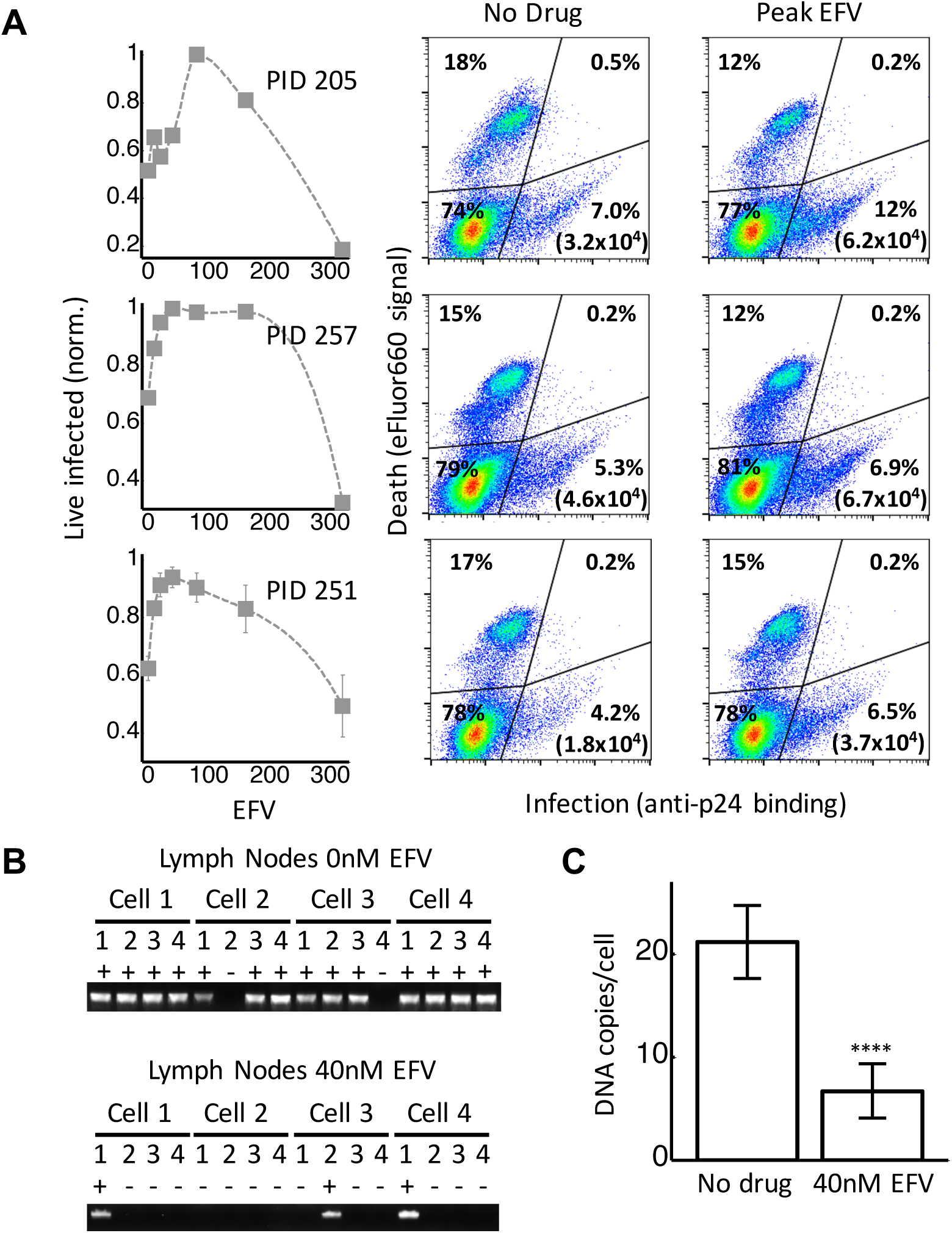
Infection optimum with partial inhibition by EFV in lymph node cells. (A) The number of live infected cells as a function of EFV. Each row represents *in vitro* infection of lymph node cells from one participant designated by a participant identification number (PID). Left column is the number of live infected cells normalized by the maximum number of live infected cells in coculture infection as a function of EFV. Middle and right columns are flow cytometry dot plots showing infection in the absence of drug and at the infection optimum, with HIV p24 on the x-axis and death as detected by the dye eFluor660 on the y-axis. Infected live cells are in the bottom right quadrant. The percent of cells in each quadrant is shown, and number in brackets represents the density of live infected cells per ml. There were sufficient cells for single experiments for lymph nodes from PID 205 and 257. Means and standard errors for 3 independent experiments are shown for the lymph node from PID 251. Dotted lines are a guide to the eye. (B) HIV DNA copy number per cell was quantified by sorting fixed p24 positive lymph node cells from PID 251 into individual wells of a multi-well plate at 1 cell per well. Cells were lysed and de-crosslinking performed. Each cell lysate was further divided into 4 wells and nested PCR performed to detect HIV DNA. First row is representative of the number of positive wells per cell for cells in the absence of drug. Second row is representative of the number of positive wells per cell for cells in the presence of 40nM EFV. (C) Quantified number of HIV DNA copies in the absence of drug and presence of 40nM EFV after correction for assay sensitivity. Results are of 56 cells from 3 independent experiments for each drug condition (n=8 for experiment 1, n=24 for experiment 2, n=24 for experiment 3). ^****^p=4x10^−9^ by 2 tailed t-test.

In each of the lymph nodes tested, we observed a peak in live infected cells at intermediate EFV concentrations. The infection optimum in the lymph node of participant 205 was visible as a plateau between 50 and 200 nM EFV (Figure 5A, first row). The lymph node from participant 251 showed a peak at 100nM EFV (Figure 5A, second row). In the presence of EFV, there was a decrease in the fraction of dead cells that was offset by a similar increase in the fraction of live infected cells for lymph nodes from both participants. There were more overall detectable cells with EFV, resulting in differences in the absolute concentrations of live infected cells being larger than the differences in the fractions of live infected cells between EFV and non-drug treated cells (Figure 5A, with absolute number of live infected cells shown in parentheses in the flow cytometry plots). This is most likely due to cells which died early becoming fragments and so being excluded from the total population in the absence of EFV.

We used a lymph node from a third individual, where we obtained more cells, to examine the number of HIV DNA copies per cell. This lymph node also showed an infection optimum at 50nM EFV (Figure 5A, third row). To detect the effect of EFV on integrations per cell, we sorted single cells based on p24 positive signal, de-crosslinked to remove the fixative (Materials and Methods), then divided each cell into 4 wells. As with the cell line, we detected HIV DNA copies by PCR 48 hours post-infection. We observed multiple DNA copies in EFV untreated lymph node cells. The number of copies decreased with EFV (Figure 5B). We corrected for sensitivity of detection as quantified in ACH-2 cells (Materials and Methods). The corrected numbers were 21 HIV DNA copies with no drug, and 5 copies in the presence of EFV at the infection optimum (Figure 5C, see Figure 5–figure supplement 1 for histograms of raw HIV DNA copy numbers per cell). Hence, the decrease in the number of copies still results in sufficient copies to infect the cell.

Since L100I does not often occur in the absence of other drug resistance mutations according to the Stanford HIV Drug Resistance Database (Rhee et al, 2003), we repeated the experiment with the K103N mutant, a frequently observed mutation in virologic failure with a higher level of resistance to EFV relative to the L100I mutant. We used cell-free infection to obtain drug inhibition per virion at each level of EFV, which we denote *d*_*103*_ (Figure 6A, see Figure 6–figure supplement 1 for logarithmic y-axis plot). The fits showed a monotonic decrease with IC_50_=26.0nM and Hill coefficient of 1.5 (Figure 6A, black line). We then proceeded to use the K103N mutant in coculture infection, using cells from two different lymph nodes in different experiments (see Figure 6–figure supplement 2 for results of individual experiments). The peak in the number of live infected cells in the presence of drug was between 80 and 160nM EFV (Figure 6B). Fits were obtained to Equation (2) using *d*_*103*_ values and the number of DNA copies in the absence of drug measured for L100I infection. The fits recapitulated the experimental results when *r* =0.7 and *q* =0.06 (Figure 6B, black line).

**Figure 6:**
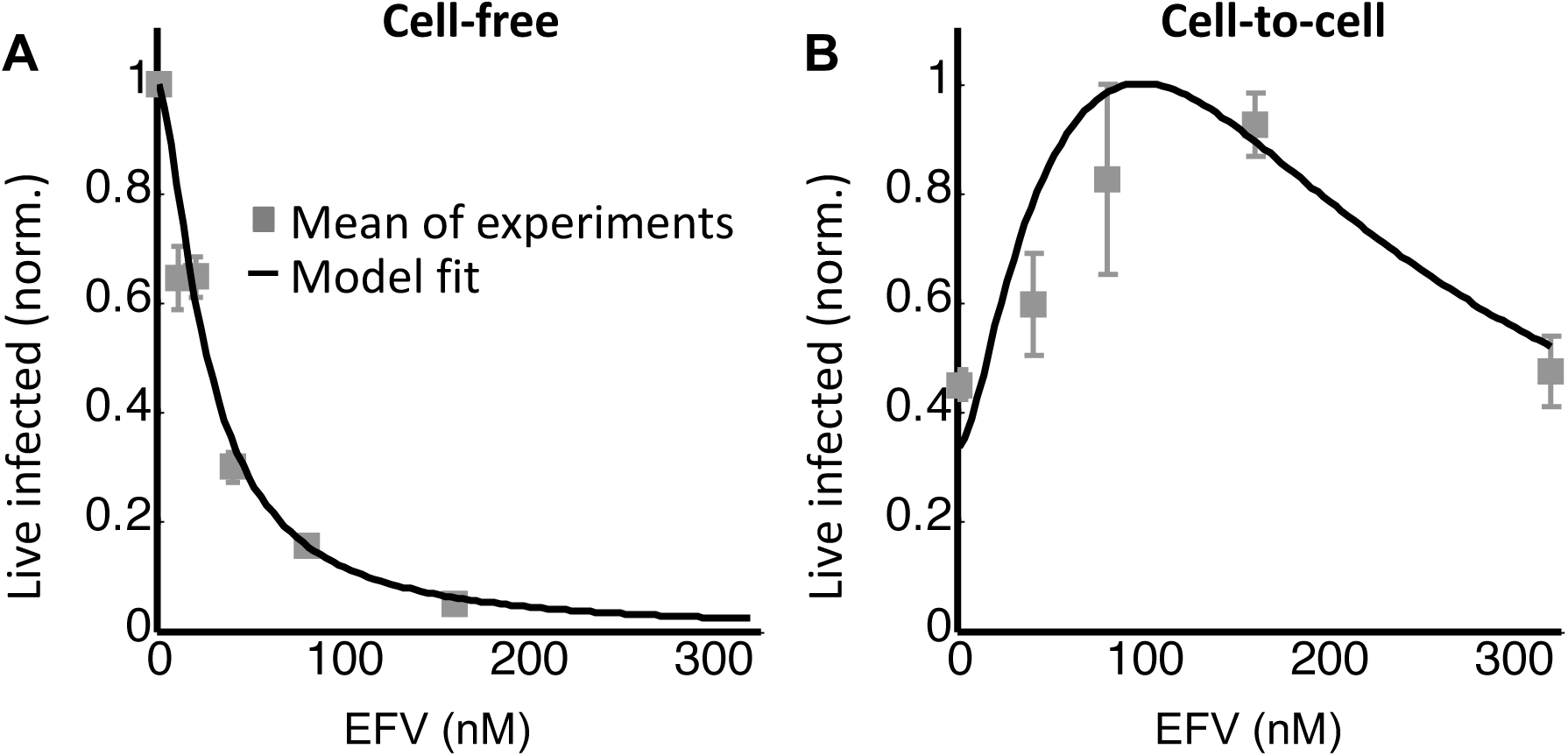
Infection with the K103N mutant shows an infection optimum at clinically observed lymph node EFV concentrations. (A) The number of live infected cells normalized by the maximum number of live infected cells in cell-free infection as a function of EFV for the K103N mutant. Black line represents best-fit model for the effect of EFV on infection with single virions, with IC_50_=26nM and Hill coefficient of 1.5. Shown are means and standard errors for 2 independent experiments using cells from PID 251. (B) The number of live infected cells normalized by the maximum number of live infected cells in coculture infection as a function of EFV for the K103N mutant. Black line represents best-fit model for the effect of EFV on infection with multiple virions according to Equation (2), with *d* values calculated based on the cell-free infection data for the K103N mutant and the mean number of integrations in the absence of drug determined for L100I. The fits recapitulated the experimental results when *r* =0.7 and *q* =0.06. Shown are means and standard errors for 3 independent experiments, There were sufficient lymph node cells from PID 251 for 2 of the 3 experiments, and the third experiment was performed with lymph node cells from PID 274.

## Discussion

The optimal virulence concept in ecology proposes that virulence needs to be balanced against host survival for optimal pathogen spread (Bonhoeffer et al, 1996; Bonhoeffer & Nowak, 1994; Gandon et al, 2001; Jensen et al, 2006). At the cellular level, this implies that the number of successfully infected cells may increase when infection virulence is reduced. The current study is, to our knowledge, the first to address this question experimentally at the level of individual cells infected with HIV.

Using a model where cells are infected and die in a probabilistic way, we found that there were two possible outcomes of partially inhibiting infection. In the case where cells were infected by single virions, inhibition always led to a decline in the number of live infected cells, since inhibition reduced the number of infections per cell from one to zero. In contrast, in the case where multiple virions infected a cell, the possibility existed that inhibition reduced the number of integrating virions, without extinguishing infection of the cell completely. If each HIV integration increases the probability of cell death, reducing the number of integrations without eliminating infection should lead to an increased probability of infected cell survival, and consequently to an increase in the number of live infected cells.

We investigated the outcome of partial inhibition of infection in both a cell line and primary lymph node cells. In both systems, we observed that there was a peak in live infected cell number at intermediate inhibitor concentrations. This correlated to a decreased number of viral DNA copies per cell. Further increasing inhibitor concentration led to a decline in live infected cell numbers, and infecting with EFV resistant mutants shifted the peak in live infected number to higher EFV concentrations. Our model as described by Equation (2) reproduced the essential behaviour of the experimental results. The model can be further refined using a distribution for the number of DNA copies per cell, and indeed the number of HIV DNA copies per lymph node cell in the presence of EFV seems bimodal (Figure 5–figure supplement 1B), with highly infected cells and cells with fewer HIV copies. Given that the simpler model was sufficient to explain the general behaviour of coculture infection in the face of inhibitor, we did not introduce a distribution in our model to avoid adding an extra unmeasured parameter.

These observations reinforce previous results showing that HIV integrations are cytotoxic (Cooper et al, 2013). In addition to HIV cytotoxicity caused by viral integrations through the mechanism of double strand breaks, other mechanisms of HIV induced death are also present, including IFI16 dependent innate immune system sensing of abortive reverse transcripts following non-productive infection of resting T cells (Doitsh et al, 2014; Monroe et al, 2014). The experiments presented here reflect the effect of partial inhibition on productive infection of HIV target cells, which mostly consist of activated T cell subsets, not resting T cells. More complex models would be needed to decipher the effect of partial inhibition of HIV infection on resting T cell numbers and the outcome of this in terms of available T cell targets in future infection cycles.

The clinical implications of an infection optimum in the presence of EFV with EFV sensitive HIV strains are likely to be negligible, since the drug concentrations at which an infection optimum occurs are extremely low. However, for EFV resistant HIV, the infection optimum shifts to the range of EFV concentrations observed in lymph nodes (~ 100nM) (Fletcher et al, 2014b), and can be expected to shift to even higher EFV concentrations with more resistant mutants. As EFV has a longer half-life than the other antiretrovirals co-formulated with it, it may be the only agent present in partially adherent individuals for substantial periods of time (Taylor et al, 2007). Therefore, partial inhibition of HIV infection with EFV may provide a surprising advantage to EFV resistant mutants, and may allow individuals failing therapy to better transmit drug resistant strains.

## Materials and Methods

### Ethical statement

Lymph nodes were obtained from the field of surgery of participants undergoing surgery for diagnostic purposes and/or complications of inflammatory lung disease. Informed consent was obtained from each participant, and the study protocol approved by the University of KwaZulu-Natal Institutional Review Board (approval BE024/09).

### Inhibitors, viruses and cells

The following reagents were obtained through the AIDS Research and Reference Reagent Program, National Institute of Allergy and Infectious Diseases, National Institutes of Health: the antiretroviral EFV; RevCEM cells from Y. Wu and J. Marsh; HIV molecular clone pNL4-3 from M. Martin. Cell-free viruses were produced by transfection of HEK293 cells with pNL4-3 using TransIT-LT1 (Mirus) or Fugene HD (Roche) transfection reagents. Virus containing supernatant was harvested after two days of incubation and filtered through a 0.45μm filter (Corning). The number of virus genomes in viral stocks was determined using the RealTime HIV-1 viral load test (Abbott Diagnostics). The L100I and K103N mutants were evolved by serial passages of wild type NL4-3 in E7 RevCEM cells in the presence of 20nM EFV. After 16 days of selection, the reverse transcriptase gene was cloned from the proviral DNA and the mutant reverse transcriptase gene was inserted into the NL4-3 molecular clone. To produce the E7 RevCEM cell clone, RevCEM cells were subcloned at single cell density and screened for the fraction of GFP expressing cells upon HIV infection using microscopy. Cell culture and experiments were performed in complete RPMI 1640 medium supplemented with L-Glutamine, sodium pyruvate, HEPES, non-essential amino acids (Lonza), and 10% heat-inactivated FBS (Hyclone).

### Infection

For a cell-free infection of E7 RevCEM and lung lymph node cells, 10^6^ cells/ml were infected with 2×10^8^ NL4-3 viral copies/ml (~20ng p24 equivalent) for 2 days. For coculture infection, infected cells from the cell-free infection were used as the donors and cocultured with target cells. For E7 RevCEM, 2% infected donor cells were added to uninfected targets and cocultured for 2 days. For lymph node cells, a ratio of 1:4 donor to targets cells was used and infection incubated for 4 days.

### Staining and flow cytometry

E7 RevCEM reporter cells were stained with 300nM of the live cell impermeable dye DRAQ7 (Biolegend) immediately before flow cytometry and live infected cells detected as the number of DRAQ7 negative, GFP positive cells. Lymph node cells were resuspended in 1ml of phosphate buffered saline (PBS) and stained at a 1:1000 dilution of the eFlour660 dye (eBioscience) according to the manufacturer’s instructions. Cells were then fixed and permeabilized using the BD Cytofix/Cytoperm Fixation/Permeabilization kit (BD Biosciences) according to the manufacturer’s instructions. Cells were then stained with anti-p24 FITC conjugated antibody (KC57, Beckman Coulter). Live infected lymph node cells were detected as the number of eFluor660 negative, p24 positive cells. Cells were acquired with a FACSAriaIII or FACSCaliber machine (BD Biosciences) using 488 and 633nm laser lines. Results were analysed with FlowJo software. For single cell sorting to detect the number of HIV DNA copies per cell, cells were single cell sorted using 85 micron nozzle in a FACSAriaIII or FACSAria Fusion machine. GFP positive, DRAQ7 negative E7 RevCEM were sorted into 96 well plates (Biorad) containing 30μl lysis buffer (2.5μl 0.1M Dithiothreitol, 5μl 5% NP40 and 22.5μl molecular biology grade water (Kurimoto et al, 2007)). Fixed, p24 positive, eFluor660 negative lymph node cells were single cell sorted into 96 well plates containing 5ul of PKD buffer (Qiagen) with 1:16 proteinase K solution (Qiagen) (Thomsen et al, 2016). Sorted plates were snap frozen and kept at -80°C until ready for PCR.

### Determination of HIV DNA copy number in individual cells

Plates were thawed at room temperature and spun down. Fixed cells were de-crosslinked by incubating in a thermocycler at 56°C for 1 hour. The lysate from each well was split equally over 10 wells (2.5μl each well after correction for evaporation) for E7 RevCEM or 4 wells (6.8μl each well after correction for evaporation) for lymph nodes, containing 50μl of Phusion hot start II DNA polymerase (New England Biolabs) PCR reaction mix (10ゼ5l 5X Phusion HF buffer, 1ゼ5l NTPs, 2.5ゼ5l of the forward primer, 2.5ゼ5l of the reverse primer, 0.5μl Phusion hot start II DNApolymerase, 2.5μl of DMSO and molecular biology grade water to 50μl reaction volume). Two rounds of PCR were performed. The first round reaction amplified a 700bp region of the reverse transcriptase gene using the forward primer 5’ CCTACACCTGTCAACATAATTGGAAG 3’ and reverse primer 5’ CATCAGAAAGAACCTCCATTC 3’. Cycling program was 98°C for 30 seconds, then 34 cycles of 98°C for 10 seconds, 63°C for 30 seconds and 72°C for 15 seconds with a final extension of 72° C for 5 minutes. 1µl of the first round product was then transferred into a PCR mix as above, with nested second round primers (forward 5’ TAAAAGCATTAGTAGAAATTTGTACAGA 3’, reverse 5’ CATCTGTTGAGGTGGGGATTTACC 3’). The second round PCR amplified a 550bp product which was then visualized on a 1% agarose gel. PCR reactions were found to work best if sorted plates were thawed no more than once, and plates which underwent repeated freeze-thaw cycles showed poor amplification.

### Correction of raw number of detected DNA copies for detection sensitivity

A stochastic simulation in Matlab was used to generate a distribution for the number of positive wells per cell for each mean number of DNA copies per cell *λ*. The probability for a DNA copy to be present within a given well and be detected was set as *σ*/*w*, where *σ* was the detection sensitivity calculated as the number of ACH-2 with detectable integrations divided by the total number of ACH-2 cells assayed (19/72, *σ* = 0.26), and *w* was the number of wells. A random number *m* representing DNA copies per cell from a Poisson distribution with a mean *λ* was drawn, and a vector *R* of *m* random numbers from a uniform distribution was generated. If there existed an element R_i_ of the vector with a value between 0 and *σ*/*w*, the first well was occupied. If an element existed with a value between *σ/w+ε* and 2(*σ/w),* the second well was occupied, and if between (*σ*/*w*+*ε*)(*n*-1) and n(*σ*/*w*), the *nth* well was occupied. The sum of wells occupied at least once was determined, and the process repeated *j* times for each *λ*, where *j* was the number of cells in the experimental data. A least squares fit was performed to select *λ* which best fit the experimental results across well frequencies, and mean and standard deviation for *λ* was derived by repeating the simulation 10 times.

### Fit of the EFV response for single infections using IC_50_ and Hill coefficient

To obtain *d*, we normalized Equation (2) by the fraction of infected cells in the absence of drug (Sigal et al, 2011) to obtain *Tx =* (infected targets with EFV)/ (infected targets no EFV)= *((1- (1-r)*^*λ/d*^*) (1-q)*^*λ/d*^*)/ ((1- (1-r)*^*λ*^ *) (1-q)*^*λ*^ ). We approximate the result at small *r, q* to *Tx =* (*1- e*^*-rλ /d*^*) e*^*-qλ /d*^*/* (*1- e*^*-rλ*^ *) e*^*-qλ*^ *= e*^*qλ (^1^-^1^/d)*^ *((1- e*^*-rλ /d*^*) / (1- e*^*-rλ*^ )). Expanding the exponentials we obtain *Tx = (1+qλ (1-1/d))* ((*-rλ/d)/-rλ) = (1+qλ (1-1/d))(1/d).* We note that at *λ < 1, qλ (1-1/d)< < 1,* and hence *Tx @ 1/d*. *Tx* was measured from the experiments to obtain *d* values at the EFV concentrations used for cell-free infection, where *λ* < 1. To obtain a fit of *d* as a function of IC_50_ and Hill coefficient (*h*) for EFV, we used the relation the relation for the fraction of unaffected infections (*F*_*u*_, (Shen et al, 2008)), whose definition is equivalent to *Tx* at *λ*<1:

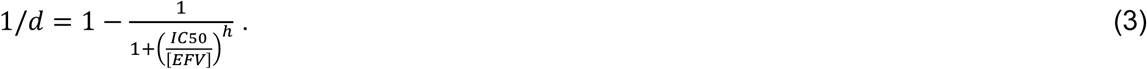

## Acknowledgements

AS was supported by a Human Frontiers Science Program Career Development Award CDA 00050/2013. RAN is supported by the European Research Council through grant Stg. 260686. LJ is supported by a fellowship from the South African National Research Foundation. IMF is supported through the Sub-Saharan African Network for TB/HIV Research Excellence (SANTHE), a DELTAS Africa Initiative (grant #DEL-15-006).

## Author contributions

AS and LJ designed the study. RM, JB, KJD, CC, YG, KK, and FK designed, coordinated and implemented collection and processing of lymph node samples. LJ, JH, SC, IF, AY, MB, and GL performed the experiments. AS and RAN derived the mathematical model. AS performed the simulations and model fits. All authors discussed the results. AS, LJ, JH, IF, and RAN wrote the manuscript.

---

**Figure 2 – Figure supplement 1.**
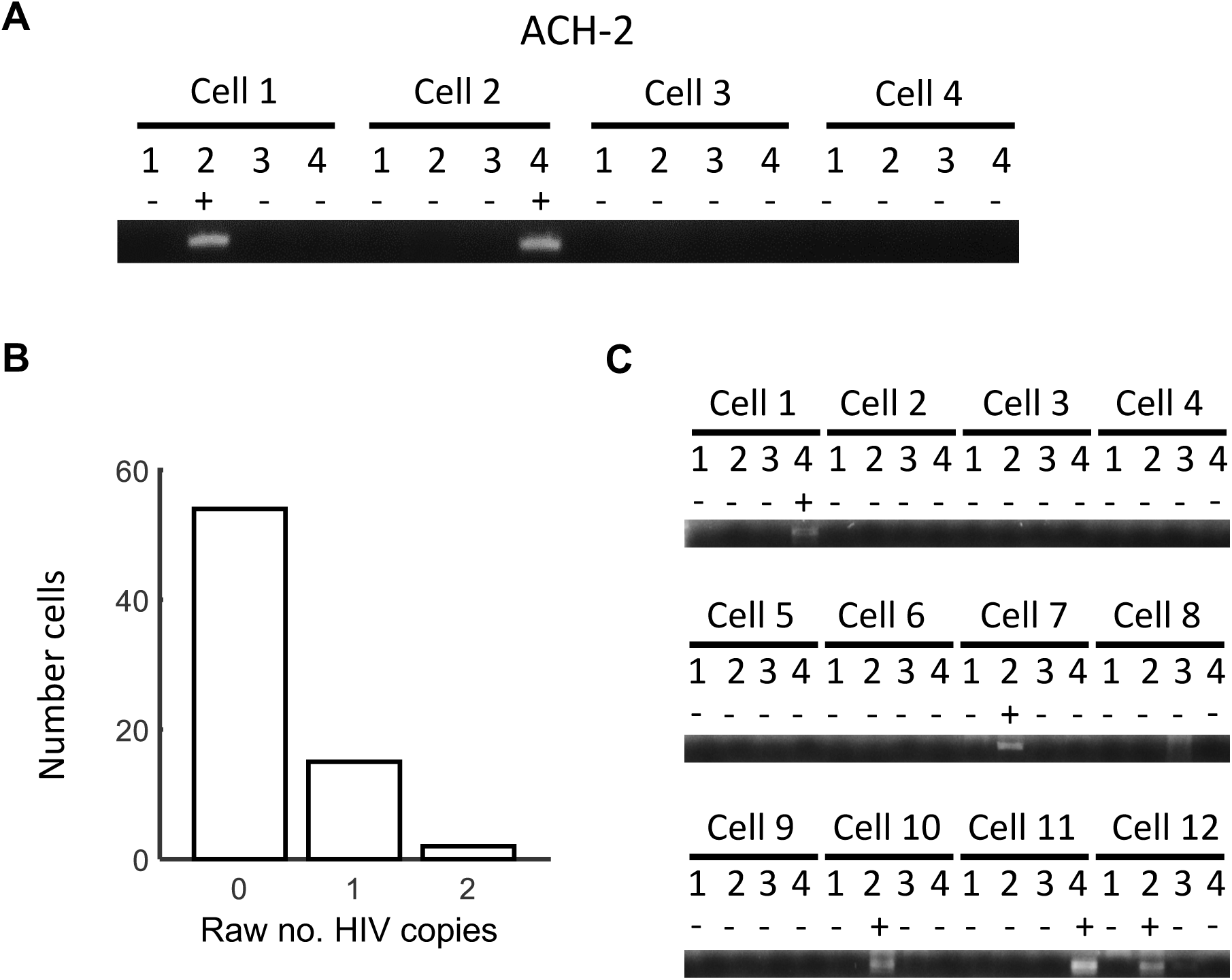
Detected integrations in ACH-2 cells and cells infected with cell-free virus. (A) To quantify HIV DNA copy number per cell in ACH-2 cells which contain single inactive integrations of HIV, cells were sorted into individual wells of a multi-well plate at 1 cell per well and lysed. Each lysed cell was further divided into 4 wells and nested PCR performed to on a region of the reverse transcriptase gene to detect whether the well contained a copy of HIV DNA. A total of 72 cells were tested in 3 independent experiments, 24 cells per experiment. (B) Histogram of the number of HIV DNA copies per cell. (C) GFP positive E7 cells infected with cell-free virus at an infection frequency of approximately 0.1 were sorted into individual wells of a multi-well plate at 1 cell per well and lysed. Each lysed cell was further divided into 4 wells and nested PCR performed. One experiment was performed, and the first 12 out of 17 cells assayed are shown.

**Figure 2 – Figure supplement 2.**
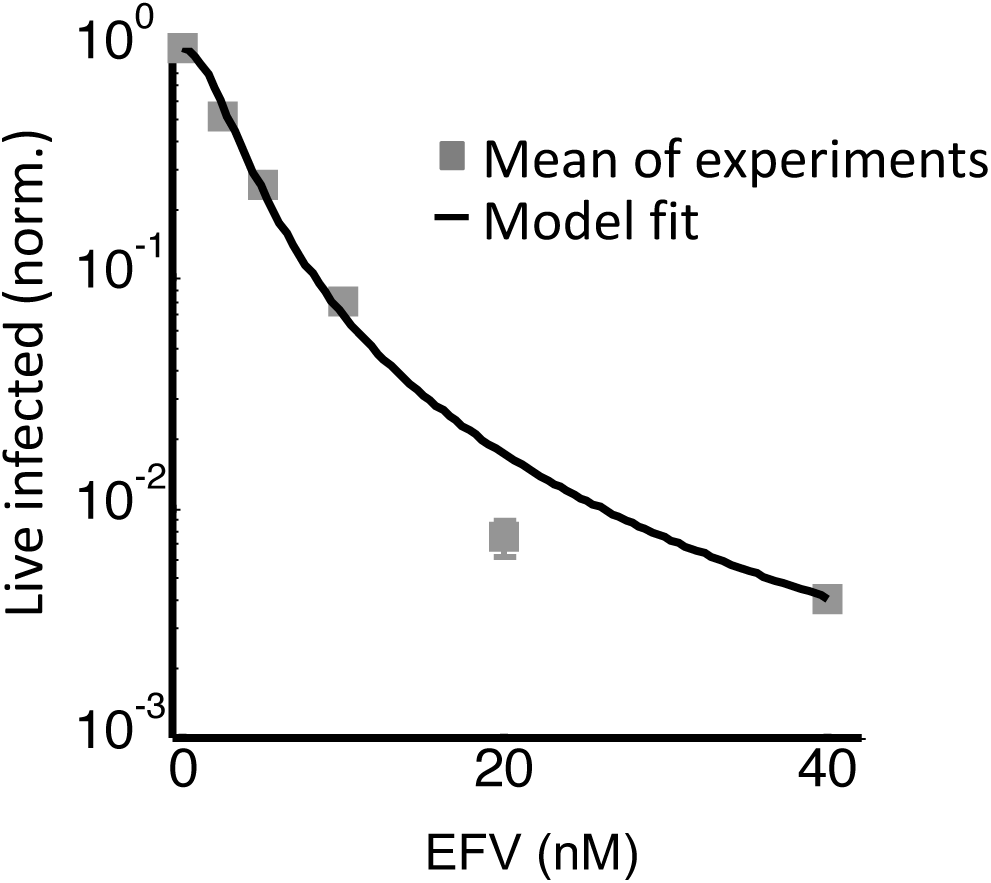
The number of live infected cells in cell-free infection with wild type HIV. Live infected cells are normalized by the maximum number of live infected cells in cell-free infection as a function of EFV. Black line represents best-fit model for the effect of EFV on infection with single virions, with IC_50_=2.4nM and Hill coefficient of 1.8. Shown are means and standard errors for 3 independent experiments. Y-axis shown on a log scale to better view fits at low live infected cell numbers.

**Figure 3 – Figure supplement 1.**
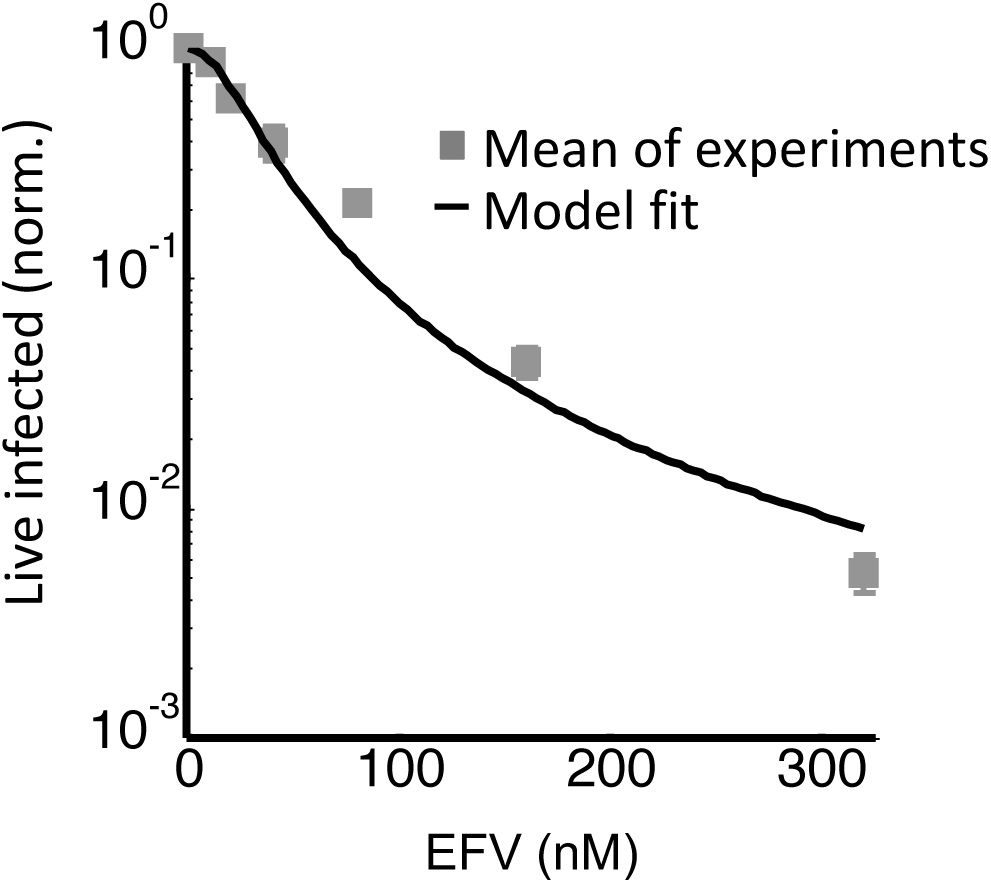
The number of live infected cells in cell-free infection with the L100I mutant. Live infected cells are normalized by the maximum number of live infected cells in cell-free infection as a function of EFV. Black line represents best-fit model for the effect of EFV on infection with single virions, with IC_50_=26.5nM and Hill coefficient of 2.0. Shown are means and standard errors for 3 independent experiments.

**Figure 5 – Figure supplement 1.**
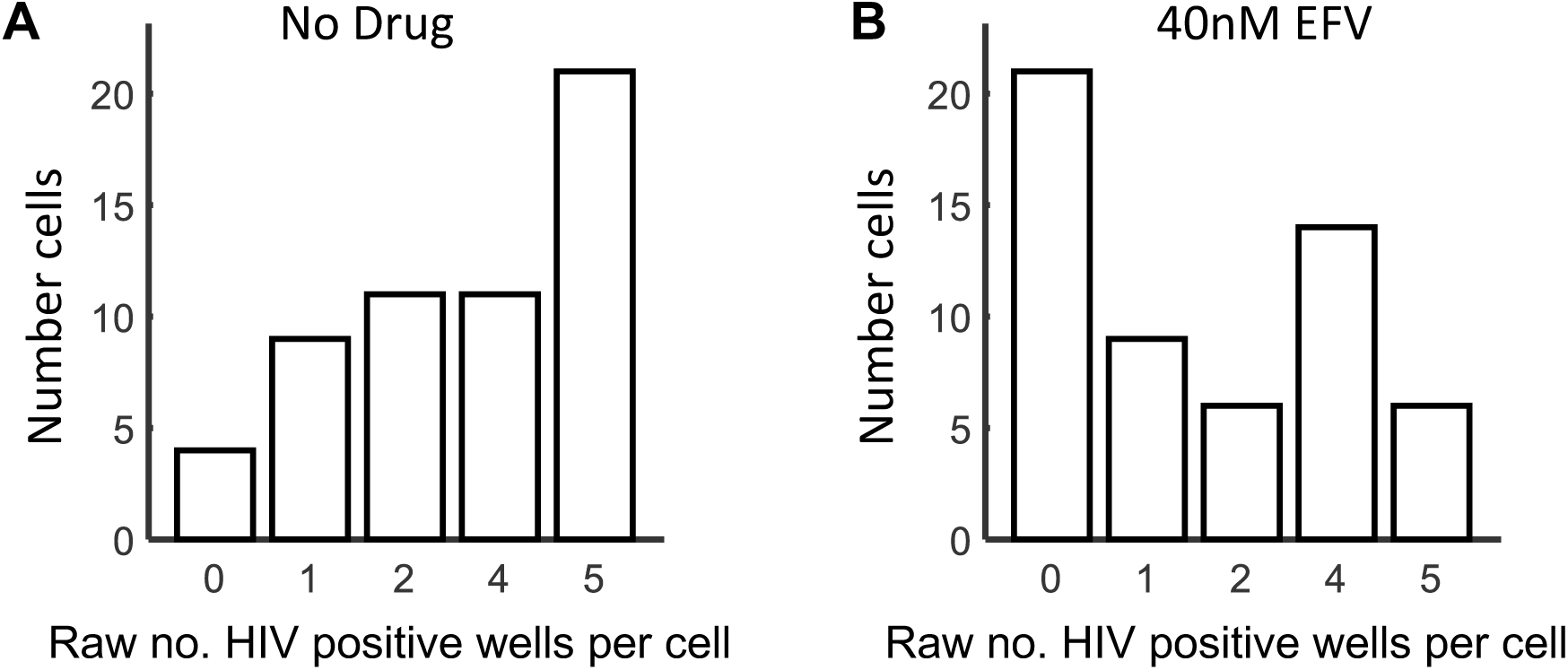
Raw HIV DNA copy numbers per lymph node cell. Fixed, p24 positive cells from PID 251 were sorted into individual wells of a multi-well plate at 1 cell per well and lysed. Each lysed cell was further divided into 4 wells and nested PCR performed on a region of the reverse transcriptase gene. Shown are histograms of the raw numbers of positive wells per cell. (A) No drug. (B) 40nM EFV. Results are from 3 independent experiments.

**Figure 6 – Figure supplement 1.**
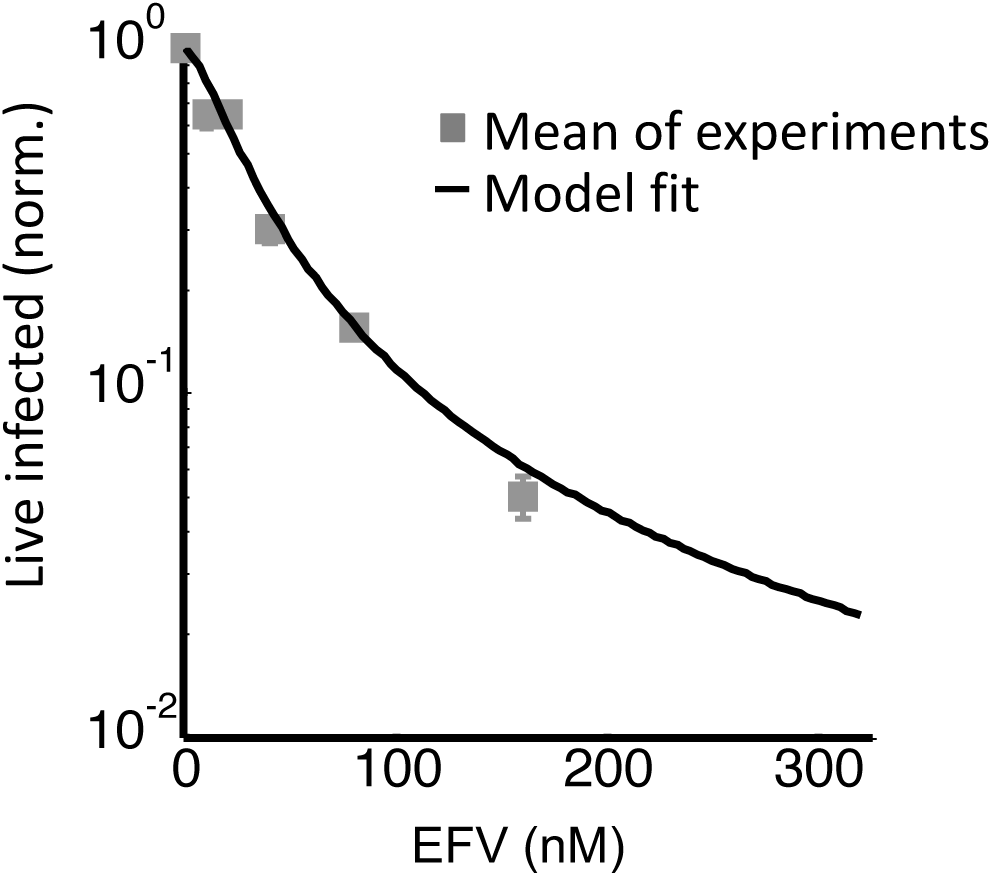
The number of live infected cells in cell-free infection with the K103N mutant. Live infected cells are normalized by the maximum number of live infected cells in cell-free infection as a function of EFV. Black line represents best-fit model for the effect of EFV on infection with single virions, with IC_50_=26.0nM and Hill coefficient of 1.5. Shown are means and standard errors for 3 independent experiments.

**Figure 6 – Figure supplement 2.**
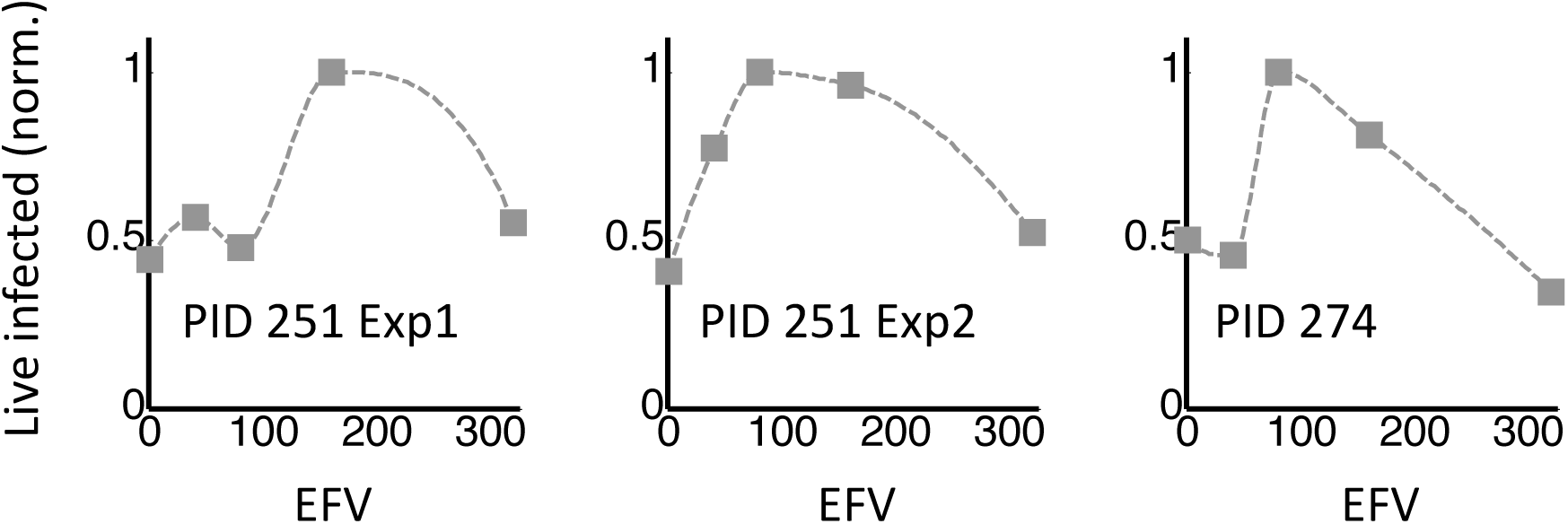
Response of K103N coculture infection to EFV in individual experiments. Shown is the number of live infected cells normalized by the maximum number of live infected cells in coculture infection as a function of EFV for the patient designated by the three digit identifier appearing below the curve. Two experiments per performed for lymph nodes resected from PID 251, and one for PID 274. Dotted lines are a guide to the eye.

**S Table 1.** Participant information.

| **Participant ID** | **Sex** | **Age (years)** | **LN location** |
| --- | --- | --- | --- |
| 205 | F | 27 | Supraclavicular |
| 251 | M | 32 | Mediastinal |
| 257 | F | 46 | Peribronchial |
| 274 | M | 22 | Peribronchial |

